# High stretchability, strength and toughness of living cells enabled by hyperelastic vimentin network

**DOI:** 10.1101/446666

**Authors:** Jiliang Hu, Yiwei Li, Yukun Hao, Tianqi Zheng, German Alberto Parada, Huayin Wu, Shaoting Lin, Shida Wang, Xuanhe Zhao, Robert D. Goldman, Shengqiang Cai, Ming Guo

**Affiliations:** Department of Mechanical Engineering, Massachusetts Institute of Technology, Cambridge, MA 02139, USA.; School of Engineering and Applied Sciences, Harvard University, Cambridge, MA 02138,USA.; Department of Cell and Molecular Biology, Northwestern University Feinberg School of Medicine, Chicago, IL 60611, USA.; Department of Mechanical and Aerospace Engineering, University of California, San Diego, La Jolla, CA 92093,USA.

## Abstract

In many normal and abnormal physiological processes, including cellular migration during normal development and invasion in cancer metastasis, cells are required to withstand severe deformations. The structural integrity of eukaryotic cells under small deformations has been known to depend on the cytoskeleton including actin filaments (F-actin), microtubules and intermediate filaments (IFs). However, it remains unclear how cells resist severe deformations since both F-actin and microtubules fluidize or disassemble under moderate strains. Here, we demonstrate that vimentin intermediate filaments (VIFs), a marker of mesenchymal cells, dominate cytoplasmic mechanics at large deformations. Our results show that cytoskeletal VIFs form a stretchable, hyperelastic network. This network works synergistically with other dissipative cytoplasmic components, substantially enhancing the strength, stretchability, resilience and toughness of the living cytoplasm.

---

Mesenchymal cells such as fibroblasts play central roles in many physiological processes including the Epithelial to Mesenchymal Transition (EMT) that takes place during normal embryonic development, in cancer metastasis and in wound healing (*1, 2*). During these physiological processes, mesenchymal cells experience severe deformations as they engage in the migratory and invasive activities associated with these processes (*3*), highlighting the importance for cells to maintain mechanical integrity under large deformations. Recent studies have emphasized the importance of nuclear envelope rupture and repair mechanisms in limiting DNA damage during mechanical stress (*4, 5*). In contrast, the mechanisms involved in protecting the remaining cytoplasm and its constituent organelles against severe deformations remain unclear.

The ability of eukaryotic cells to resist deformation depends on the cytoskeleton, an interconnected network of biopolymers including F-actin, MT and IFs (*6*). Recent work has demonstrated that both F-actin and MT structures fluidize at moderate strains (20% and 60%, respectively) (*7*), suggesting that they cannot maintain the mechanical integrity and resilience of the cytoplasm at even larger strains, which can happen in the physiological processes mentioned above. Therefore, it has been hypothesized that cytoplasmic IFs may play an important role in maintaining the mechanical integrity and resilience of cells, especially under large deformations (*8*). As a key phenotypic marker of mesenchymal cells, vimentin IFs (VIFs) are known to be critical for regulating cell shape, migration (*9*), and cytoplasmic stiffness at small deformations (*10, 11*). However, their structural and mechanical roles in living mesenchymal cells at large deformations remain elusive and unclear.

Here we show that VIFs behave as a strain-stiffening hyperelastic network in mesenchymal cells and that they determine cellular strength, stretchability, resilience and toughness. VIF networks interconnect with other cytoskeletal networks and effectively disperse local deformations in the cytoplasm, and thus lead to a significant increase of the mechanical energy input required to damage the cytoplasm. Furthermore, the hyperelastic VIF network can dramatically slow down both poroelastic relaxation and viscoelastic relaxation processes in the cytoplasm, which can enhance the mechanical damping capability of the cytoplasm and thus provide better protection for the interior organelles. Our results provide a fundamental insight into the essential role of VIFs for maintaining cell structural and mechanical integrity and resilience during a variety of key physiological processes.

To study the effect of VIFs in cytoplasmic mechanics, we use wild type (WT) and vimentin null (Vim^-/-^) mouse embryonic fibroblasts (mEFs); these cells have been shown to have the normal arrays of other cytoskeletal systems as described in previous studies (*10, 12*). To demonstrate the importance of VIFs in cells undergoing large deformations, we encapsulate living WT and Vim^-/-^ mEFs in biocompatible hydrogels composed of alginate and polyethylene glycol (PEG). The PEG- alginate hydrogel is both tough and highly-stretchable; therefore mEFs encapsulated in this 3D matrix can be highly deformed by stretching the hydrogel (*13*). Hydrogels with encapsulated mEFs are subjected to 5 different stretching regimes ranging from 0% to 300% and cell viability is determined by carrying out cell viability (live/dead) assays (Fig. 1A and Fig. S1). We find that WT mEFs maintain a high viability (∼ 90%) under stretch, while the viability of Vim^-/-^ mEFs decreases significantly as the strain increases (Fig. 1B). This result reveals the key role of VIFs in maintaining cell viability under large deformation.

**Fig. 1.**
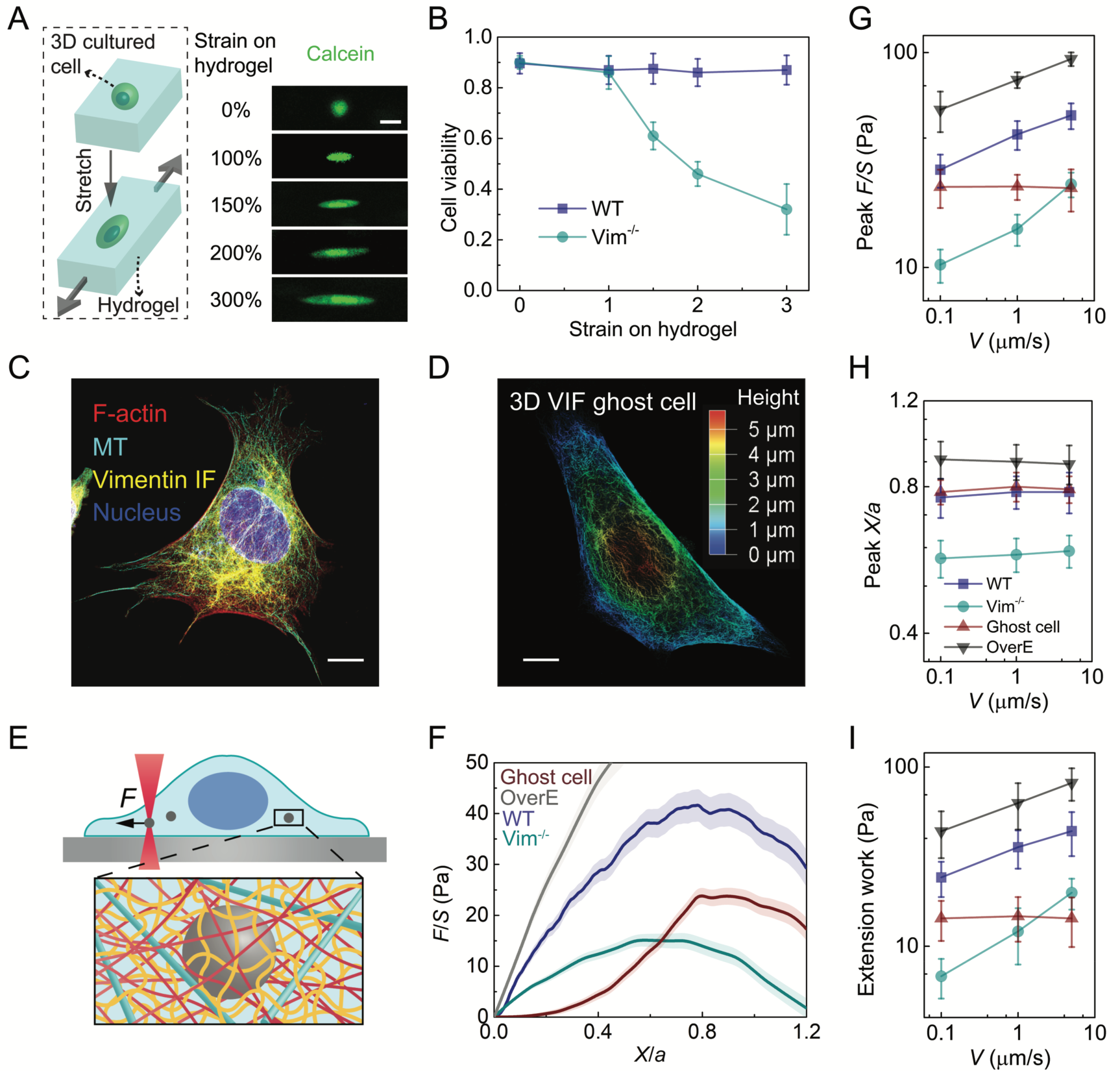
VIF network maintains cell viability under large deformations and increases the mechanical strength, stretchability and toughness of the cytoplasm. **(A)** Cells cultured in 3D PEG-alginate hydrogels are deformed by stretching the hydrogels. Cells are labeled with calcein to visualize cell shape and viability (scale bar: 20 μm). **(B)** The viability of WT and Vim^-/-^ mEFs are measured with live/dead assay under various strains for 2 hours (n=15). **(C)** Immunofluorescence image of WT mEF showing the microtubule and vimentin IF cytoskeletons (scale bar: 10 μm). **(D)** 3D microscopic image of VIF in a ghost cell; the color represents height (scale bar: 10 μm). **(E)** Schematic of micro-mechanical measurements in the cytoplasm using optical tweezers. **(F)** Normalized force-displacement curves obtained in cells at *V* = 1μm/s, from which we calculate peak *F/S*, peak *X/a* and extension work. The semi-transparent band around the average curves represents the standard error (n=20 cells for each curve). **(G - I)** The dependence of peak *F/S* (G), peak *X/a* (H) and extension work (I) on loading rate in different cells. Error bars represent standard deviation (n=20 cells).

To understand these findings, we study how VIFs impact the structural integrity and mechanical behavior of living cells. To accomplish this, polystyrene beads (diameter *a* = 1 μm) are introduced into living WT and Vim^-/-^ mEFs through endocytosis. These beads are much larger than the typical cytoskeleton mesh size (∼ 50 nm) and thus can probe the cytoplasm as a continuous medium (*14*). To probe cytoplasmic mechanics, we use optical tweezers to drag a bead unidirectionally toward the nucleus with a constant speed of 1.0 μm/s, as illustrated in Fig. 1E (*15*). The resultant resistant force (*F*) from the surrounding cytoplasmic structures continuously increases with the displacement (*X*) of the bead until a peak force is reached, after which the material yields (Fig. 1F). The normalized peak force (peak *F/S,* where *S* is the bead cross-sectional area) and the normalized displacement under peak *F/S* (defined as peak *X/a*) are determined in order to characterize the cytoplasmic strength and stretchability, respectively (Figs. 1G and 1H); both peak *F/S* and peak *X/a* are significantly larger in WT mEFs than in Vim^-/-^ mEFs, showing that VIFs substantially increase the cytoplasmic strength and stretchability. Moreover, we calculate the extension work by integrating the normalized force-displacement curve (from *X/a*=0 to *X/a*=1.2) to characterize cytoplasmic toughness. The extension work of WT mEFs (35.7 ± 8.6 Pa) is about 3 times that of Vim^-/-^ mEFs (12.1 ± 4.2 Pa), which indicates that VIFs can significantly improve cytoplasmic toughness.

To further investigate the mechanical properties of cytoplasmic VIF networks, major cellular components including cell membranes, F-actin and microtubules are extracted from WT mEFs (*16*) while leaving only the VIF network structure *in situ* as a “ghost cell” (Fig. S2). The VIF network structures seen in live cells are preserved in the ghost cells, as shown in reconstituted 3D images (Fig. 1D and Movie S1). Under small deformations, the resistant force measured by dragging a bead in a ghost cell is lower than that in Vim^-/-^ mEFs at the same displacement (Fig. 1F). However, the force required to further deform the ghost cell exceeds that of Vim^-/-^ mEFs due to the strong strain stiffening behavior of the VIF networks (*17*); surprisingly, it reaches a peak *F/S* of 23.8 ± 3.2 Pa, which is markedly larger than that of Vim^-/-^ cells (15.1 ± 2.5 Pa). Indeed, the ghost cell has a similar peak strain (*X/a*) as that of WT mEFs (Fig. 1H). These results show that VIFs determine the mechanical behavior of the cytoplasm under moderate to severe deformations. Furthermore, as we overexpress VIFs in WT mEFs (OverE mEFs), we find that that the strength, stretchability and toughness of the cytoplasm are further enhanced (Fig. 1G-I and Fig. S3). This critical role of VIFs in enhancing the strength, stretchability and toughness of the cytoplasm is very important for a variety of physiological processes that involve large cellular deformations, such as those seen in embryonic development and in metastasis.

Cells are additionally known to behave as viscoelastic and poroelastic materials whose mechanical behavior strongly depends on the deformation rate (*15, 18, 19*). To further characterize the impact of loading rate on the role of VIFs, force-displacement relationships are determined in WT mEFs, Vim^-/-^ mEFs and VIF ghost cells under a range of loading speeds (0.1 to 5.0 μm/s), as shown in Fig. S4. Interestingly, we find that the peak *F/S*, peak *X/a* and the extension work are independent of loading speed in the ghost cell (Fig. 1G-I), suggesting that VIFs can be regarded as a purely hyperelastic network. In contrast, the peak *F/S* and extension work increase with loading speed in both WT and Vim^-/-^ mEFs (Fig. 1H, I), reflecting the combined rate-dependent nature of the remaining other cytoskeletal components.

To further study the effect of VIFs on cytoplasmic rate-dependent behavior, relaxation tests are carried out in cells by applying an instant deformation (*X/a* = 0.4) with a 1 μm-diameter bead using optical tweezers. We then hold the bead and record the corresponding resistant force as a function of time. The resistant force in the VIF ghost cell slightly relaxes (relaxed *F/S* = 0.75 ± 0.40 Pa) at short timescale (*t* < 0.05 s), and remains at a steady plateau over the experimental timescales employed (0.05s < *t* < 10 s) (Fig. 2A-C). The initial force relaxation in the VIF ghost cell at short timescales follows an exponential decay (inset of Fig. 2C), which corresponds to poroelastic relaxation (*15, 19*). The poroelastic effect in the cytoplasm accounts for the resistance to cytosol flow through the porous cytoskeletal structure. Poroelastic fitting (*F/S* ∼ exp(-*t/t*_p_)) of the averaged relaxation curve in the VIF ghost cell yields a poroelastic relaxation time tp ≈ 0.028s, which is consistent with previous measurements in mammalian cells (*15, 19*). This result suggests that VIF networks act as a porous and hyperelastic meshwork exhibiting no viscous effects, which is consistent with macroscopic rheology tests on reconstituted VIF networks (*7*). In WT and Vim^-/-^ mEFs, the force relaxation curves exhibit an exponential decay followed by a power law decay (Fig. 2C). The initial exponential decay regime and the long-time power law decay regime correspond to poroelastic and viscoelastic relaxations of living cells, respectively (*18, 19*). Taken together, the results further reveal that the VIF network is hyperelastic but the other cytoplasmic components are viscoelastic, while all of them share an intrinsically porous nature.

**Fig. 2.**
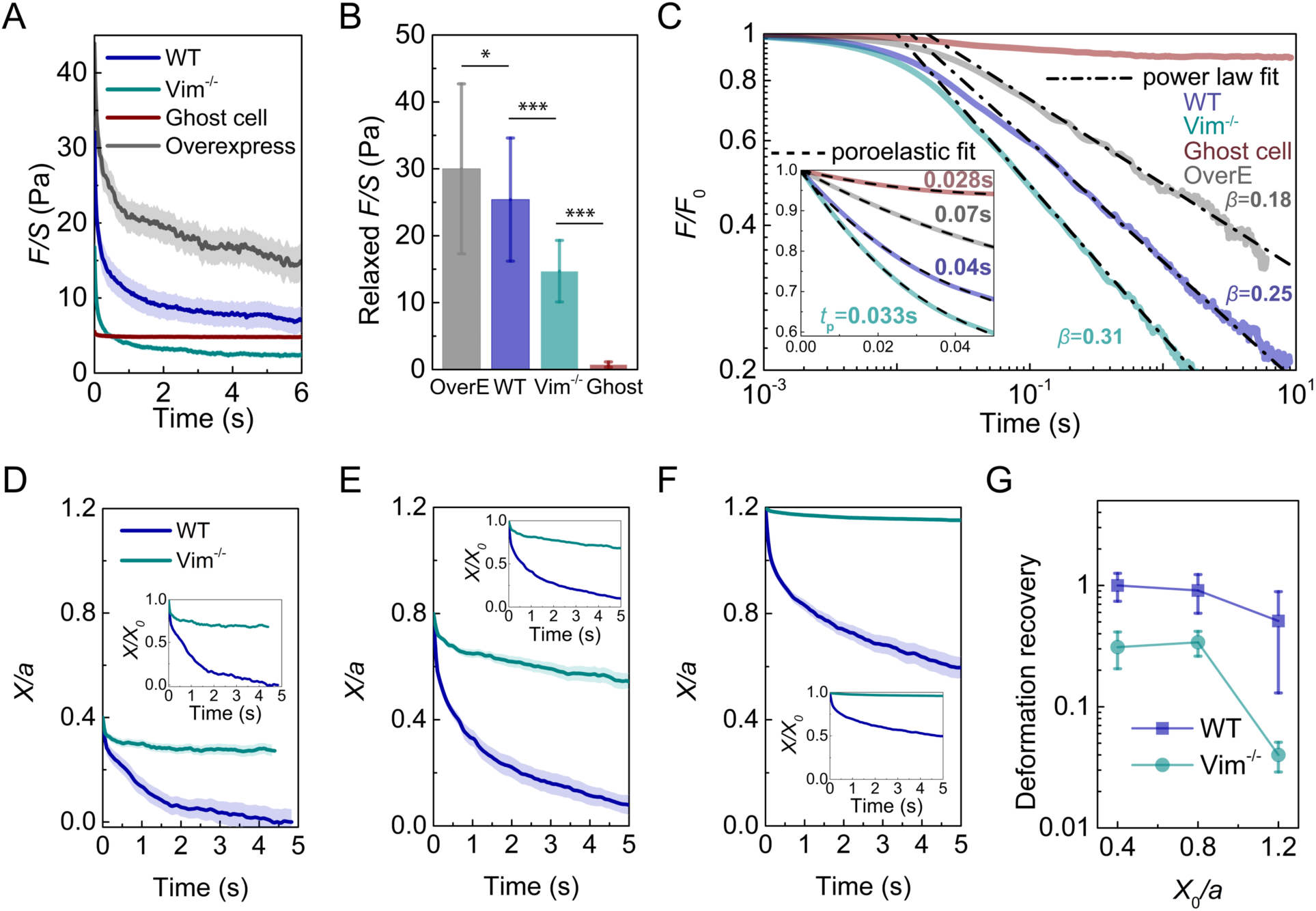
The elastic VIF network increases the relaxed force, relaxation time and yielding strain of the cytoplasm. **(A)** Force relaxation curves of different cells under an instant initial *X/a* = 0.4. The semi-transparent band around the average curves represents the standard error (n=15 cells for WT and overexpress, n=25 cells for Vim^-/-^ and ghost cell). **(B)** Comparison of cellular relaxed force, which is defined as the decrease of *F/S* over the relaxation test. Error bars represent standard deviation (n=15 cells for WT and overexpress, n=25 cells for Vim^-/-^ and ghost cell). **(C)** Relaxation curves are normalized with initial *F*_0_ at *t* = 0. The curves are fitted with viscoelastic power law decay at long time scales (0.05s < *t* < 10s), and are fitted with poroelastic exponential decay (inset) at short time scales (*t* < 0.05s). **(D - F)** Deformation recovery in the cytoplasm of WT and Vim^-/-^ mEFs under different initial normalized displacement (*X*_0_/*a* = 0.4, *X*_0_/*a* = 0.8, *X*_0_/*a* = 1.2, respectively). The semi-transparent band around the average curves represents the standard error (n=15 cells for each curve). **(G)** The cytoplasmic deformation recovery (defined as the recovered deformation over initial deformation) under different initial deformations in WT and Vim^-/-^ mEFs.

The relaxed normalized force (relaxed *F/S*) is higher in the WT mEFs (25.4 ± 9.2 Pa) than that in the Vim^-/-^ mEFs (14.7 ± 4.6 Pa), as shown in Fig. 2B. Interestingly, the presence of VIFs markedly increases the poroelastic relaxation time in cells; we find that t_p_ ≈ 0.04s in the WT mEFs, and 0.033s in the Vim^-/-^ mEFs (Fig. 2C, inset). This suggests that VIFs reduce the average cytoplasmic mesh size. Moreover, as we fit the relaxation curves at long time scales (*t* > 0.1s, as shown in Fig. 2C) to a power law function, *F/S* ∼ *t^-β^,* to characterize the viscoelastic relaxation speed of the cytoplasm, we find a lower relaxation speed (*β* ≈ 0.25) in the WT mEFs compared to the Vim^-/-^ mEFs (*β* ≈ 0.31). This might be due to friction and other interactions between VIF networks and other cytoskeletal components. Consistently, both the relaxation time and the relaxed force further increase in the OverE mEFs (Fig. 2C). The increased relaxed force and relaxation times in the WT and OverE mEFs, compared with the Vim^-/-^ mEFs, imply that VIF networks can regulate the mechanical damping capacity of the cytoplasm, and thereby provide an optimal protection for organelles while maintaining the structural integrity of cells.

The relaxation test results indicate that VIF networks remain elastic up to deformations of *X/a* = 0.4. To study the yielding strain (the strain limit after which the material exhibits a plastic response) of VIF networks in living cells, we apply different deformations (*X/a* = 0.4 – 1.2) by dragging a 1 μm-diameter bead at 1μm/s using optical tweezers. After reaching the expected initial displacement, we release the force applied on the bead by turning off the laser power, subsequently recording the movement of the released bead by microscopic imaging. After releasing the loading force, the bead moves backward with time (Fig. 2D–F), indicating an elastic recovery of the cytoplasmic deformation. We find that there is full recovery of deformations up to *X/a* = 0.8 in WT mEFs, while the Vim^-/-^ mEFs begin to exhibit plastic deformation (i.e. not fully recovered) for deformations *X/a* below 0.4. This result suggests that VIF networks can increase the yielding strain and thus the resilience of the cytoplasm, providing living cells with a mechanism for recovering their original shapes and structures after large deformations.

The capacity of energy absorption is an important parameter characterizing materials; it is defined as the material’s toughness, and can be obtained by integrating the full stress-strain curve until fracture (*20*). Here, we integrate the normalized force-displacement curve to *X/a*=1.2, and obtain the extension work of cells (Fig. 1I). We find that the extension work is larger in the WT mEFs than the sum of those in the Vim^-/-^ mEFs and the VIF ghost cell (Fig. S4). Since we locally deform the cytoplasm using a bead, complete fracture of the cytoplasm is not achieved at the end of the loading; nevertheless, this comparison suggests that the cytoplasmic toughness of the WT mEFs is greater than the superposition of the Vim^-/-^ mEFs and the VIF ghost cell, indicating a direct interaction between VIFs and the rest of the cytoplasmic components.

To further investigate the underlying mechanisms by which VIFs enhance the toughness of the cytoplasm, cyclic loading and unloading tests are carried out in cells (inset of Fig. 3A). This test quantifies both the elastic and dissipated mechanical energy that together constitute material toughness. Surprisingly, the loading and unloading curves obtained in the VIF ghost cell collapse, and remain unchanged over more than 100 cycles (Fig. 3A). This result further demonstrates that cytoplasmic VIF network itself is hyperelastic and does not dissipate energy (Fig. 3D). Although individual VIF have been shown to dissipate mechanical energy under large strains due to the unfolding of α helices (*21, 22*), our results suggest that individual VIF filaments retain an elastic regime even though VIF networks withstand large deformations. When this measurement is made in the WT mEFs, a clear hysteresis loop is observed, suggesting energy is being dissipated during this process (Fig. 3B and Fig. S5). Furthermore, the cytoplasm is greatly softened over a few cycles and eventually becomes similar to that measured in the ghost cell after 10 cycles (Fig. 2B). In contrast, the resistant force in the Vim^-/-^ mEFs eventually becomes very weak (Fig. 3C), suggesting that cyclic loading causes most cytoskeletal structures to be damaged, fluidized or rearranged in cells without VIFs. After repeated cyclic loading, the normalized force-displacement curve reaches a steady state at the 10^th^ cycle in both WT and Vim^-/-^ mEFs; integration of the steady state force-displacement curve represents the elastic energy of the cytoplasm. These results show that elastic energy is significantly higher in the WT mEFs as compared to the Vim^-/-^ mEFs (Fig. 3D), further suggesting that the VIF network maintains the resilience of the cytoplasm during cyclic loading, while the rest of the cytoplasmic components contribute to the energy dissipation. To further quantify the energy dissipation, we integrate the area looped by the first loading and unloading curves (Fig. 3A-C). We find that the dissipated energy density is significantly higher in the WT mEFs than in the Vim^-/-^ mEFs (Fig. 3D), demonstrating that the existence of the VIF network can also substantially increase the energy dissipated by the cytoplasm. Since the VIF network itself is hyperelastic and does not dissipate mechanical energy (Fig. 3D), the observed enhancement in energy dissipation is more likely to be due to other cellular components through their interactions with VIFs. Similar to the design principles in interpenetrating double network tough hydrogels (*20*), the stretchy elastic VIF network and the other dissipative cytoplasmic components work synergistically, enhancing cytoplasmic toughness. Indeed, VIF network interpenetrates with other cytoskeletal networks as shown by our microscopic imaging in the cell (Fig. 1C) and previous literature (*23, 24*), providing an unavoidable physical interaction between different networks.

**Fig. 3.**
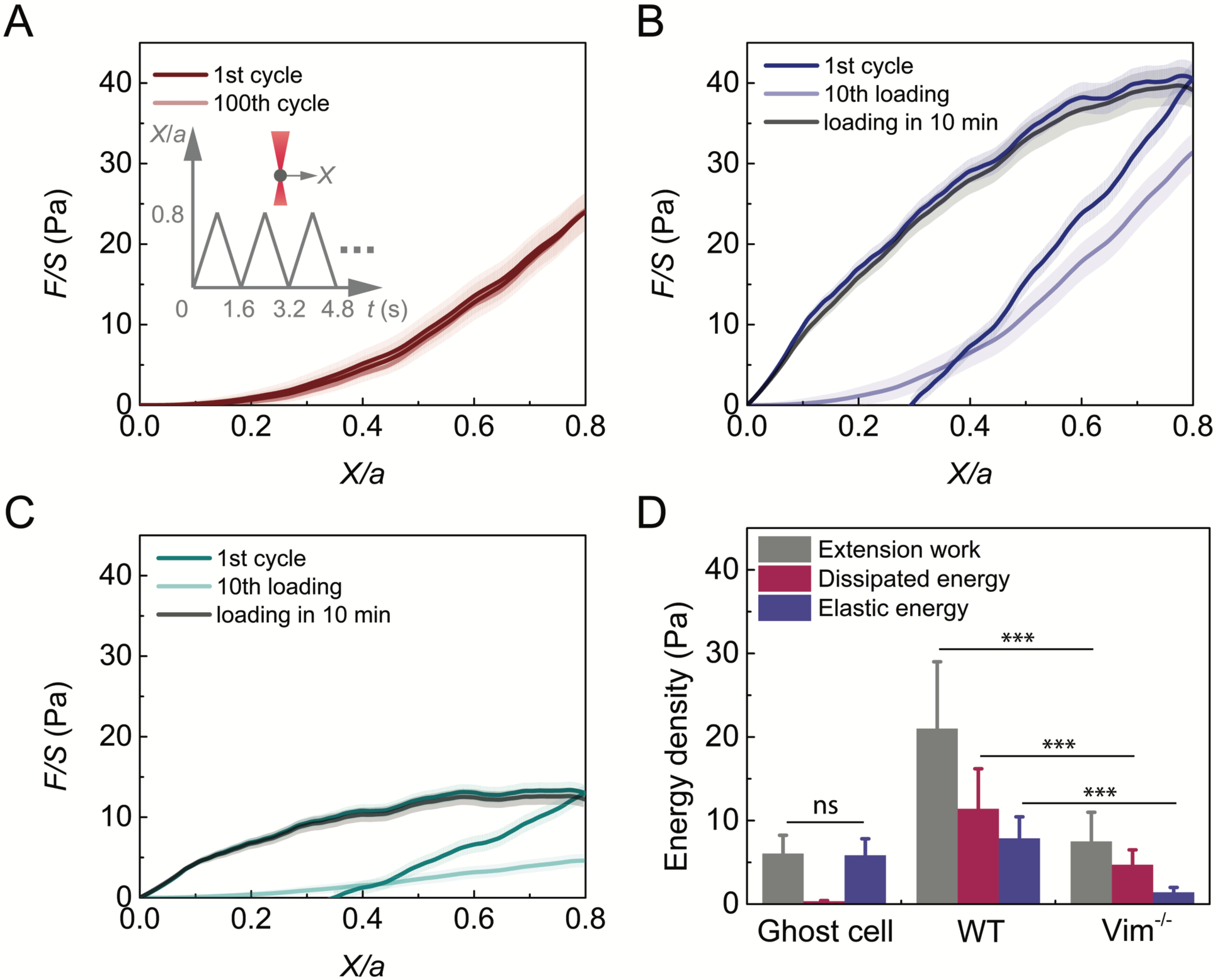
The VIF network increases the toughness of the cytoplasm by increasing dissipated energy and elastic energy. **(A)** Cyclic loading in the cell is achieved by reciprocating movement of a bead at a speed of 1 μm/s using optical tweezers (inset). The 1^st^ and 100^th^ loading and unloading curves in VIF ghost cells overlap. The semi-transparent band around the average curves represents the standard error (n=15 cells). **(B, C)** Plot of the 1^st^ and 10^th^ cyclic loading curves in WT mEFs (B) and Vim^-/-^ mEFs (C). After being damaged by repeated loadings, the loading curves in both WT and Vim^-/-^ mEFs recover to original levels in 10 minutes, highlighting the self-healing nature of the cytoplasm. The semi-transparent band around the average curves represents the standard error (n=20 cells for each curve). **(D)** VIF ghost cell does not dissipate energy. WT mEFs have significantly higher extension work, dissipated energy and elastic energy than Vim^-/-^ mEFs. Error bars represent standard deviation (n=15 cells for ghost cells, n=20 cells for WT and Vim^-/-^ mEFs).

Robust mechanical properties are essential for the survival of mesenchymal cells during cyclic deformations, since these cells are often exposed to repeated loading and large deformations (e.g. when migrating through tissues and vessels). Our results show that VIFs are essential for the cytoplasm to retain high strength, stretchability and toughness under frequent large deformations, thus preventing the cytoplasm from becoming irreversibly damaged during important cell migratory and invasive activities. Interestingly, we also find that the living cytoplasm shows significant softening after repeated loading, but can recover to its original state after a 10-minute rest (Fig. 3B, C). This highlights the quick self-reorganization ability of other cytoskeletal structures, as also observed in both mechanical testing and microscopic imaging of reconstituted actin and microtubule networks (*25-28*), even though these structures are easily disassociated upon initial loading. Such self-healing observed in the living cell cytoplasm is substantially more efficient than in typical artificial materials (*29*).

To understand the mechanisms by which VIFs enhance and regulate the mechanical properties of cytoplasm and increase the dissipated mechanical energy of the cytoplasm, we image the resultant displacement and strain fields upon introducing a local deformation in the cytoplasm. We drag a 2-micron-diameter bead in the cytoplasm over 200 nm at a speed of 2 μm/s, and perform particle image velocimetry (PIV) by visualizing 2D projected movements of surrounding fluorescently labeled mitochondria (Fig. 4A, 4B). We find that the deformation field extends significantly further in the WT mEFs than in the Vim^-/-^ mEFs (Fig. 4C, 4D, Movies S2 and S3). To quantify this effect, we plot the local cytoplasmic displacement as a function of the distance to the loading center along the drag direction (white dashed lines in Fig. 4C, 4D), where the displacement (*U*) is normalized by the bead displacement (*U*_0_), and the distance (*X*) is normalized by the bead radius (*R*), as shown in Fig. 4G. The normalized displacement decays with normalized distance as a power law with a power of −1 in the cytoplasm of the WT mEFs. In contrast, the observed displacement decay is markedly faster in the Vim^-/-^ mEFs (Fig. 4G) with a power of −2. Consistently, the normal strain field computed from the 2D displacement map concentrates around the loading point in the Vim^-/-^ mEFs while extends significantly further in the WT mEFs (Fig. 4E, 4F and Fig. S5). Taken together, these results demonstrate the contribution of VIF network in propagating the local deformation to a greater range in the cytoplasm, therefore involving more cytoplasmic materials to respond to local loading.

**Fig. 4.**
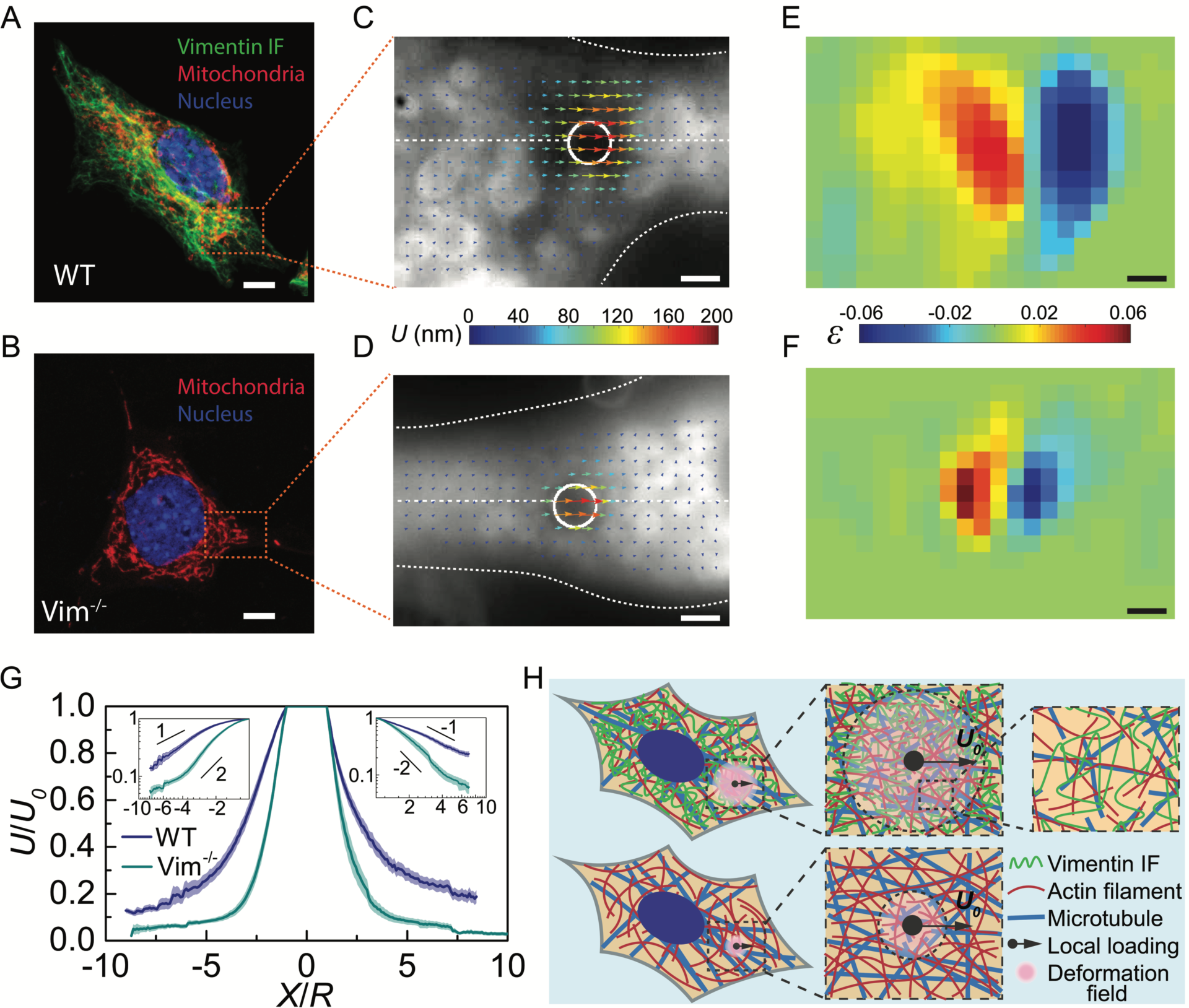
VIFs extend the cytoplasmic deformation field under local loading, acting as a stretchable hyperelastic network. **(A, B)** Deformation field is obtained by dragging a 2 μm- diameter bead in the cytoplasm over 200 nm and visualizing the 2D projected movements of surrounding fluorescently labeled mitochondria. **(C, D)** The displacement fields around the loading beads (white circle) in WT and Vim^-/-^ mEFs (cell boundaries are marked with white dotted lines). The color and the length of arrows represent displacement magnitudes. **(E, F)** The normal strain fields in WT and Vim^-/-^ mEFs are obtained from displacement fields shown in panel (C) and (D). The color represents the magnitude of normal strain. **(G)** Plot of the normalized cytoplasmic displacement as a function of the normalized distance to the loading center, along the drag directions (horizontal white dashed lines in panels (C) and (D)). Log-log plots are shown in the insets. The semi-transparent band around the average curves represents the standard error (n=15 cells for each curve). **(H)** Schematics to illustrate the mechanism of extending deformation fields by VIF network. Highly stretchable VIF networks maintain the elasticity of and transmit local strain through a large zone in the cytoplasm, while other cytoskeletal structures are easy to be damaged under deformation. Scale bars equal to 10 μm in A and B, equal to 2 μm in C – F.

Other cytoskeletal components such as F-actin and microtubules are easily relaxed and reorganized under deformation, as shown by both our measurement in cells (Fig. 2A) and macroscopic rheology measurements of reconstituted networks (*17*); once relaxed and reorganized, they quickly lose the ability to transmit stress and strain. However, the high stretchability and hyperelastic nature of the VIF network allow it to maintain structural and mechanical integrity under very large strains (more than 300%), therefore enabling local strain and stress to be effectively transmitted from a stress concentrated area to a large zone in the cell (Fig. 4H). As widely observed in common materials, stress/strain concentrations are the major cause of material damage (*30*). Under a concentrated loading, the stretchy, hyperelastic VIF network not only stores substantial mechanical energy by itself, but also increases the dissipated mechanical energy significantly by deforming larger volumes of other cellular components near the stress concentration. Therefore, the presence of such an interconnected VIF network can markedly increase the required mechanical energy to damage the cytoplasm under the same loading force.

This work uncovers the essential role of VIFs in cell mechanical behavior under large deformations. As a highly stretchable hyperelastic network, the cytoskeletal VIF network greatly enhances several of the most important mechanical properties of the cytoplasm: stretchability, resilience, strength and toughness (Table S1). The cytoplasmic stretchability is solely determined by the VIF network. Subjected to repeated loadings, the VIF network plays an essential role in maintaining the resilience of the cytoplasm because of its high yielding strain, while the rest of cytoplasmic components are greatly softened or even fluidized. By interacting with other cytoskeletal systems and organelles, VIF networks significantly enhance cytoplasmic mechanical strength and toughness under dynamic deformations, through slowing poroelastic and viscoelastic relaxations. Moreover, the stretchy VIF network can effectively propagate local stress and strain into a larger region of the cell, deforming more microtubules and actin filaments which interpenetrate and interact with the VIF network. Thus VIFs significantly enhance the strength and toughness of the cytoplasm, reducing the risk of cell damage during processes involving large deformations. The remarkable enhancement of mechanical properties by VIFs is required for mesenchymal cells to perform many physiological activities including cell migration during embryonic development, wound healing and cancer metastasis. These properties of the VIF networks may also shed light on the roles of other types of intermediate filament networks, as well as on the pathogenesis of the many human diseases associated with mutations in IF-encoding genes (*31, 32*). Furthermore, the stretchy and hyperelastic VIF network forms a dynamic yet mechanically robust cytoskeleton when combined with the dissipative and quickly recovering cytoskeletal components such as microtubules and actin filaments; such a collaborative design principle may inspire improved engineering of smart materials.

## Acknowledgements

We would like to thank P. Ronceray and the Guo Laboratory for helpful discussions. M. G. would like to acknowledge the support from the Department of Mechanical Engineering at MIT. S.C. acknowledges the support from Hellman Fellows Fund. H. W. and R. G. acknowledge the support from the National Institutes of Health (2P01GM096971-06A1).

## Data and materials availability

All data is available in the main text or the supplementary materials. All data, code, and materials used in the analysis are available upon request from the corresponding author.

**Supplementary Materials:**

Materials and Methods

Figures S1-S5

Tables S1

Movies S1-S3

References (33-39)

